# Asymmetry of auditory-motor speech processing is determined by language experience

**DOI:** 10.1101/2020.06.05.137067

**Authors:** Ding-lan Tang, Riikka Möttönen, Salomi S. Asaridou, Kate E. Watkins

**Author notes:** Correspondence: Ding-lan Tang.

## Abstract

Speech processing relies on interactions between auditory and motor systems and is asymmetrically organized in the human brain. The left auditory system is specialized for processing of phonemes, whereas the right is specialized for processing of pitch changes in speech that affect prosody. In speakers of tonal languages, however, processing of pitch (i.e., tone) changes that alter word meaning is left-lateralized. This indicates that linguistic function and language experience shape auditory speech processing asymmetries; their effect on auditory-motor speech processing remains unknown, however. Here, we investigated the asymmetry of motor contributions to auditory speech processing in speakers of tonal and non-tonal languages. We temporarily disrupted the left or right speech motor cortex using repetitive transcranial magnetic stimulation (rTMS) and measured the impact of these disruptions on auditory processing of phoneme and tone changes in sequences of syllables using electroencephalography (EEG). We found that disruption of the speech motor cortex in the left, but not the right hemisphere, impaired processing of phoneme changes in both language groups equally. In contrast, the effect of motor disruptions on processing of tone changes differed in tonal and non-tonal language groups: disruption of the left speech motor cortex significantly impaired processing of tone changes in tonal language speakers, whereas disruption of the right speech motor cortex modulated processing of tone changes in non-tonal speakers. We conclude that the contribution of the left and right speech motor cortex to auditory speech processing is determined by the functional role of the acoustic cues in the listener’s native language.

**Significance Statement:** The principles underlying hemispheric asymmetries of auditory speech processing remain debated. The asymmetry of auditory speech processing is affected by the low-level acoustic cues, but also by their linguistic function. By combining TMS and EEG, we investigated the asymmetry of motor contributions to auditory speech processing in tonal and non-tonal language speakers. For the first time, we provide causal evidence that auditory-motor speech processing asymmetries are shaped by the functional role of the acoustic cues in the listener’s native language. The lateralised top-down motor influences are likely to affect asymmetry of speech processing in the auditory system.

## Introduction

Over the past two decades, a growing number of studies have emphasized the importance of interactions between auditory and motor systems for speech perception (1-4). The dorsal pathway connecting auditory and motor systems is strongly left-lateralized for phonemic processing (5). In contrast, auditory-motor processing of prosody is lateralized to the right hemisphere (6-8).

Asymmetry in auditory processing is thought to reflect differences in the temporal and spectral sensitivities of the left and right auditory cortex to acoustic inputs (9-12). This extensive body of work indicates that the left auditory cortex is specialized for processing rapidly changing temporal information, which is necessary to resolve phonemic changes requiring discrimination of small acoustic differences in speech sounds that alter the meanings of words. In contrast, the right auditory cortex is sensitive to more slowly changing auditory information and the spectral content of the same acoustic signals, which is necessary to discriminate prosodic changes in speech, such as the use of pitch for stress, or to convey emotion. Although this view has gained lots of support, it should be noted that is has been primarily tested in speakers of non-tonal languages.

For as many as 70% of the world’s languages, supra-segmental speech elements such as changes in pitch (i.e. tones) are also used to alter word meaning (13). For example, in Mandarin Chinese, the word “ma” produced with a high-level tone (ma1) means “mother” and when produced with a rising-falling tone (ma3) means “horse”. Processing of these lexical tones in tonal language speakers is also left lateralised (14-17) but does not require rapid temporal processing of the acoustic signal. In this case, the left lateralization reflects the linguistic function of the acoustic signals and thereby demonstrates how language experience can alter hemispheric lateralization of speech processing.

It is likely that asymmetry in speech processing is determined by bottom-up properties in the acoustic signal as well as by top-down influences (11,18). We hypothesised that one such influence could be the speech motor system. We previously showed that the left speech motor cortex contributes to auditory speech processing of phonemes in non-tonal language speakers (19,20) even when there is no task to perform and processing is automatic or outside of attention (21,22). Less is known about the contribution of the right speech motor cortex to auditory speech processing and the asymmetry of such motor contributions.

Here, we aimed to determine whether speech processing in the auditory system is affected by lateralized interactions with the motor system. We investigated the asymmetry of auditory-motor speech processing in both tonal and non-tonal language groups capitalizing on the fact that the same low-level acoustic signal (e.g. pitch) serves different linguistic functions in each. Testing speakers from tonal and non-tonal language groups also allowed us to explore the influence of language experience on the motor contribution to tone processing.

Specifically, we examined the causal contributions of left and right speech motor cortex on automatic brain responses to tone and phoneme changes in speech sound sequences in tonal language (Mandarin) speakers and non-tonal language speakers. We used low-frequency repetitive transcranial magnetic stimulation (TMS) to temporarily disrupt the speech motor cortex and electroencephalography (EEG) to record mismatch negativity (MMN) responses to changes in a sequence of frequent “da” syllables (da1) that were either a phoneme (ba1) or a tone change (da4).

## Results

### Effects of TMS-induced disruption in the left speech motor cortex

We stimulated the left speech motor cortex in 15 non-tonal language speakers and 16 tonal language (Mandarin) speakers. Both groups showed significant grand-averaged MMN responses at FCz for infrequent tone (‘da4’ vs. ‘da1’) and phoneme (‘ba1’ vs. ‘da1’) changes (see Fig. 1A, Table 1 and Table S1 for group latencies and mean amplitudes).

**Table 1.** Left hemisphere stimulation: MMN peak latencies (ms) and mean amplitudes (SEM; μV) at FCz.

| No-TMS |  |  | Post-TMS |  |
| --- | --- | --- | --- | --- |
|  | Latency | Amplitude | Latency | Amplitude |
| Non-tonal language group, n=15 |  |  |  |  |
| ‘ba1’ | 182.8 | -2.97 (0.32) | 190.7 | -2.19 (0.27) |
| ‘da4’ | 189.7 | -5.80 (0.59) | 192.0 | -4.86 (0.70) |
| Tonal language group, n=16 |  |  |  |  |
| ‘ba1’ | 181.5 | -2.63 (0.30) | 199.4 | -1.75 (0.32) |
| ‘da4’ | 191.6 | -6.61 (0.70) | 187.6 | -4.91 (0.66) |

Note: The MMN amplitudes were calculated by averaging the responses across a 40-ms window centred at this peak latency.

**Figure 1.**
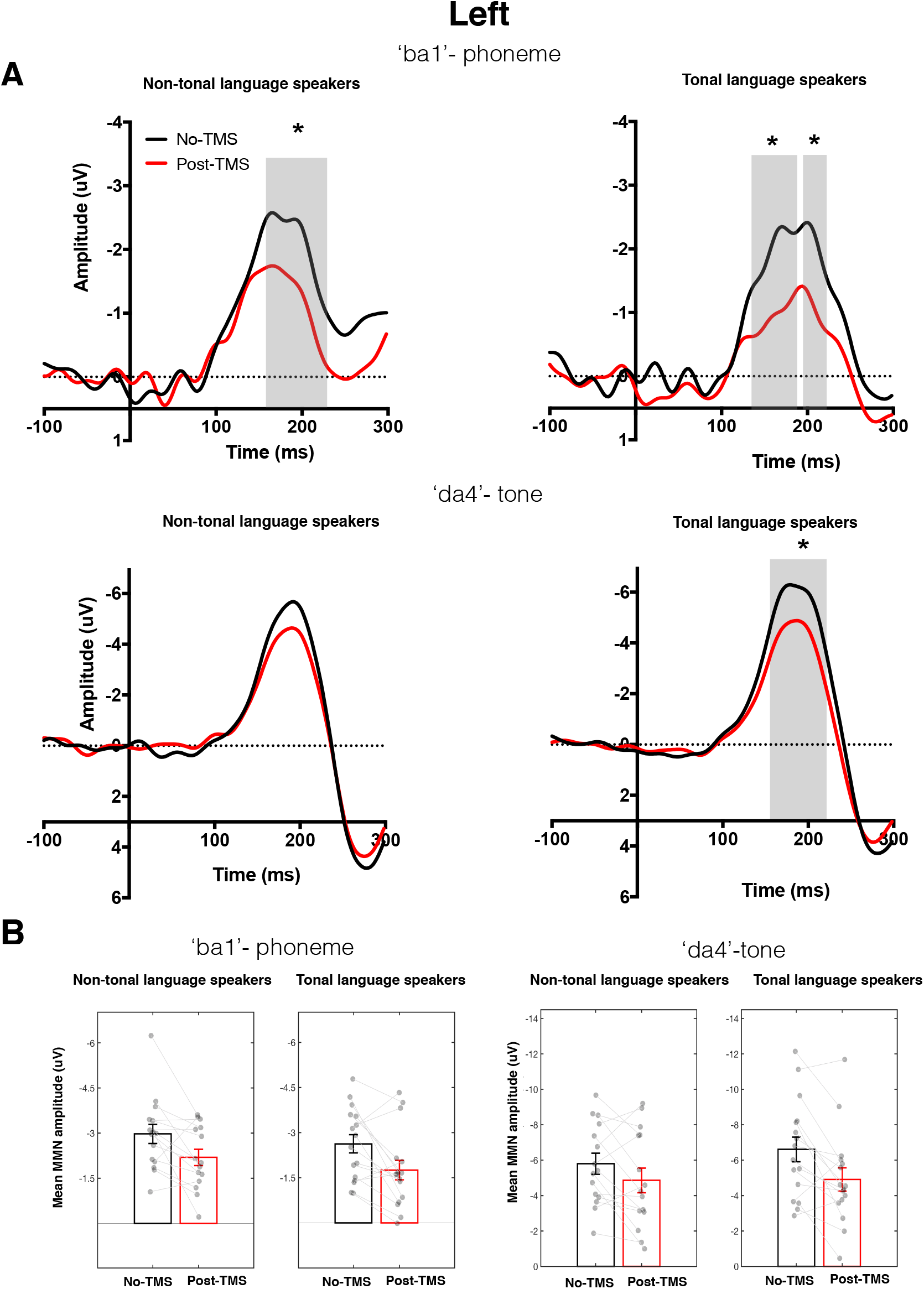
Effect of rTMS over left speech motor cortex on MMN responses elicited by the phoneme and tone changes in tonal and non-tonal language speakers. (A) The MMN responses recorded from the FCz electrode were obtained by subtracting the responses to frequent ‘da1’ syllable from the responses to infrequent ‘ba1’ (upper panel) or ‘da4’ (bottom panel) syllables. The shaded area indicates the time windows in which no-TMS (black) and post-TMS (red) MMN responses differed significantly from each other (sequential t-tests). *\*p* < .05 (2-tailed). (B) Comparison of amplitudes of MMN responses. The grey circles show the mean amplitudes of MMN responses for each participant. Mean MMN amplitudes obtained during no-TMS and post-TMS recordings for individual participants are connected by a grey line. The group mean MMN amplitudes are shown in black or red unfilled bars during no-TMS recording and post-rTMS recording respectively. Error bars represent standard error of the mean. Note the different scales used for the two stimuli; MMN responses to the tone change were robustly higher than for the phoneme change.

In the non-tonal language group, TMS disruption of left speech motor cortex significantly suppressed the amplitude of MMN responses elicited by the phoneme change 162–224 ms after stimulus onset (‘ba1’) but did not significantly modulate MMN responses to the tone change (‘da4’). In contrast, in the tonal language group, disruption of left speech motor cortex significantly suppressed the amplitude of MMN responses elicited by the phoneme change 134–190 ms and 196-224 after stimulus onset (‘ba1’) and the tone change 154–224 ms after stimulus onset (‘da4’).

Comparing the MMN responses elicited by the phoneme change (‘ba1’ vs. ‘da1’) between the two language groups, we found that the effects of left-hemisphere stimulation were not different. Both groups showed the suppressive effect of left-hemisphere stimulation and the magnitude of this effect did not differ between them (main effect of TMS: F_1,29_ = 15.56, p < .001; interaction: F_1,29_ < 1; see Fig. 1B). In contrast, the MMN responses elicited by the tone change (‘da4’ vs. ‘da1’) were supressed by left-hemisphere stimulation to a numerically larger extent in the tonal language group (mean MMN amplitude change ± SE = 1.70 μV ± 0.50) compared with the non-tonal language group (mean MMN amplitude change ± SE = 0.94 μV ± 0.50; see bottom panel of Fig. 1B) but this group difference was also not statistically significant (main effect of TMS: F_1,29_ = 13.80, p = .001; interaction: F_1,29_ = 1.14, p= .294).

In sum, our results showed that TMS-induced disruption of the left speech motor cortex suppressed MMN responses elicited by a phoneme change in both language groups. In contrast, TMS-induced disruption of the left motor cortex suppressed MMN responses elicited by a tone change in the tonal language group but brought about a nonsignificant reduction in the non-tonal language group (see Fig. 1A). It should be noted, however, that this small reduction in the MMN responses to tone discrimination during disruption of the left speech motor cortex in the non-tonal language group was statistically indistinguishable from the significant and large effect seen in the tonal language group (i.e. the interaction between group and TMS was not significant, p>.05). Examination of data shown in Fig. 1B indicates that some individuals in the non-tonal language group showed large reductions in the MMN amplitude elicited by the tone change (‘da4’) but this was not a reliable effect across the group.

### Effects of TMS-induced disruption in the right speech motor cortex

We stimulated the right speech motor cortex in another 15 non-tonal language speakers and a further 16 tonal language (Mandarin) speakers. Both groups showed significant grand-averaged MMN responses at FCz elicited by infrequent tone (‘da4’ vs. ‘da1’) and phoneme (‘ba1’ vs. ‘da1’) changes (see Fig. 2A, Table 2 and Table S2 for group latencies and mean amplitudes).

**Table 2.**
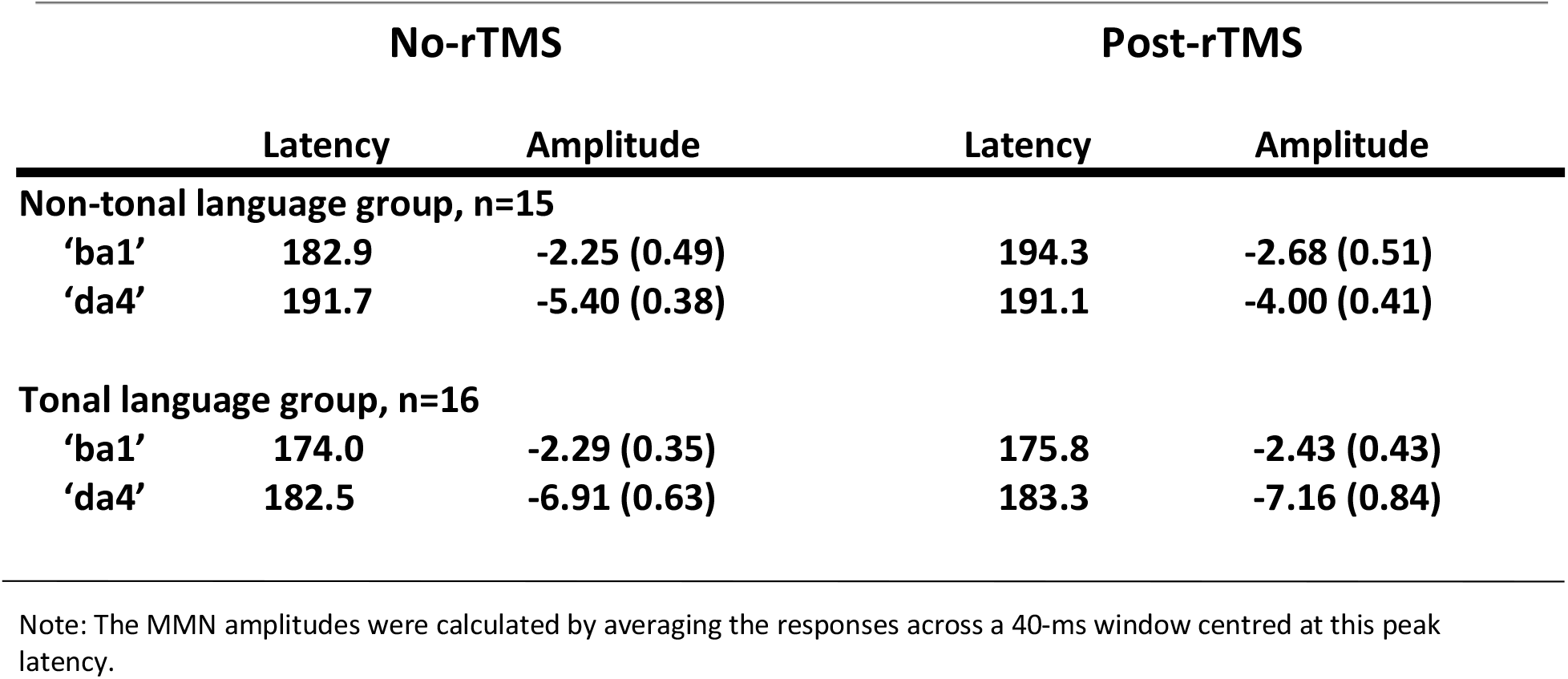
Right hemisphere stimulation: MMN peak latencies (ms) and mean amplitudes (SEM, μV) at FCz

| No-rTMS |  |  | Post-rTMS |  |
| --- | --- | --- | --- | --- |
|  | Latency | Amplitude | Latency | Amplitude |
| Non-tonal language group, n=15 |  |  |  |  |
| ‘ba1’ | 182.9 | -2.25 (0.49) | 194.3 | -2.68 (0.51) |
| ‘da4’ | 191.7 | -5.40 (0.38) | 191.1 | -4.00 (0.41) |
| Tonal language group, n=16 |  |  |  |  |
| ‘ba1’ | 174.0 | -2.29 (0.35) | 175.8 | -2.43 (0.43) |
| ‘da4’ | 182.5 | -6.91 (0.63) | 183.3 | -7.16 (0.84) |

Note: The MMN amplitudes were calculated by averaging the responses across a 40-ms window centred at this peak latency.

**Figure 2.**
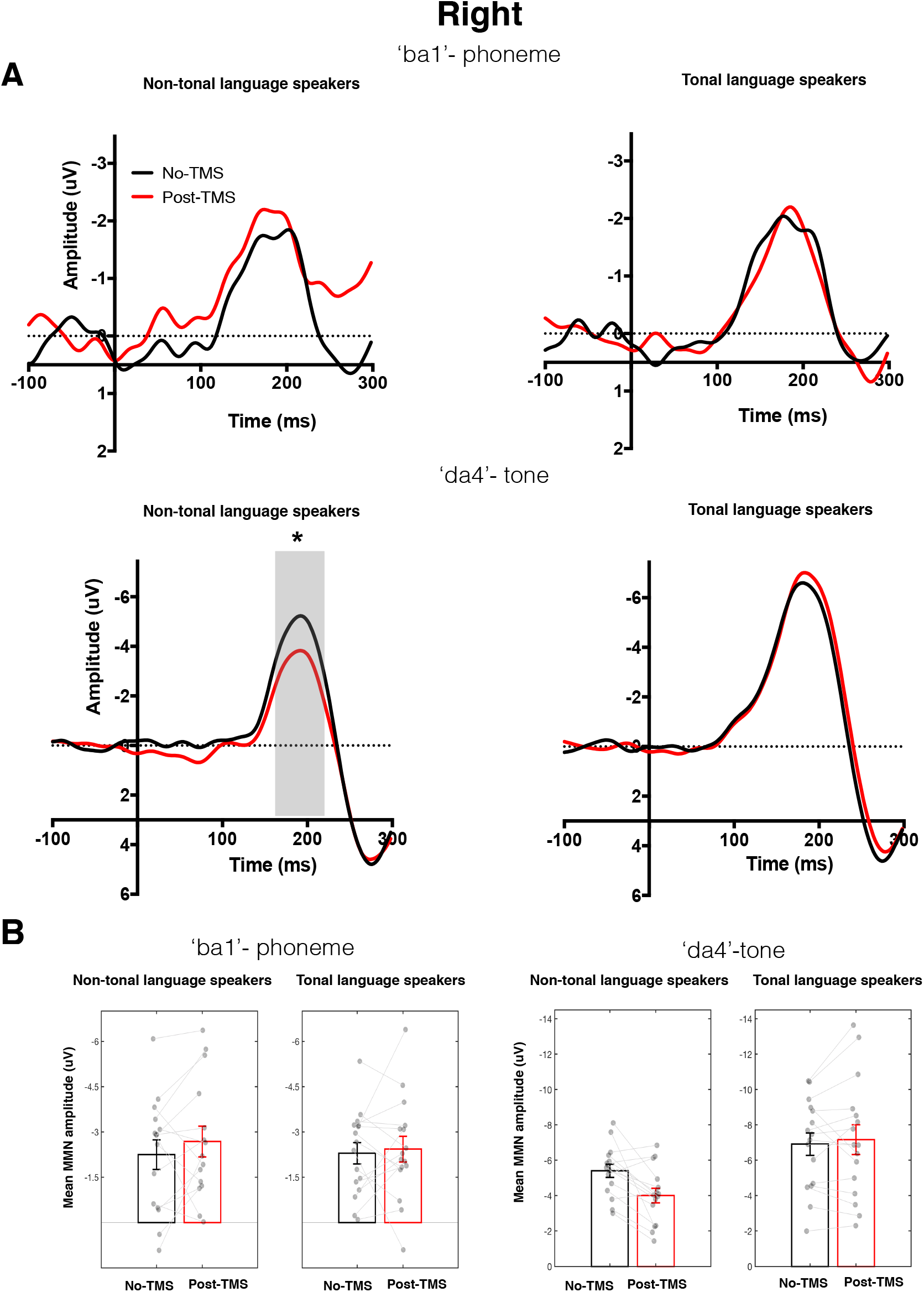
Effect of rTMS over right speech motor cortex on MMN responses elicited by the phoneme and tone changes in tonal and non-tonal language speakers. See legend to Fig. 1 for details.

In the tonal language group, TMS disruption of right speech motor cortex did not significantly modulate MMN responses elicited by either the phoneme change (‘ba1’) or the tone change (‘da4’). In contrast, in the non-tonal language group, the same TMS over the right speech motor cortex resulted in a different pattern: it had no effect on MMNs elicited by the phoneme change (‘ba1’), but significantly suppressed the amplitude of MMN responses elicited by the tone change 158–230 ms after stimulus onset (‘da4’).

For the MMN responses elicited by the phoneme change, TMS-induced disruption in the right hemisphere had no effect in either language group (i.e. the main effect of TMS was not significant: F_1,29_ = 1.02, p = .322). However, right hemisphere stimulation did affect the MMN responses elicited by the tone change differently according to language experience: it significantly suppressed these responses in the non-tonal (t_14_ = 3.41, p = .004) but not the tonal language group (t_14_ = .65, p = .526) and this group difference was significant (interaction between TMS and group: F_1,29_ = 8.57, p = .007) (Fig. 2B).

In sum, our results showed that in both language groups, TMS-induced disruption of the left but not the right speech motor cortex significantly suppressed MMN responses elicited by a phoneme change, replicating our earlier findings (21). In contrast, the effect of the right-hemisphere disruption on MMN responses elicited by a tone change differed between language groups: it had no effect on MMN responses in the tonal language group but significantly suppressed MMN responses in the non-tonal language group.

## Discussion

We used EEG and TMS to investigate the asymmetry of auditory-motor processing of phoneme and tone changes in auditory syllables in speakers of tonal and non-tonal languages. The phoneme changes have similar functional roles in tonal and non-tonal languages, whereas tone changes by definition have different functional roles. We found that disruption of the left, but not the right speech motor cortex impaired auditory discrimination of phonemes in both tonal and non-tonal language groups equally. In contrast, disruption of the right motor cortex impaired automatic auditory discrimination of tones in non-tonal language group, whereas disruption of the left motor cortex modulated automatic auditory discrimination of tones in tonal language group (i.e., Mandarin). That is, the lateralization of auditory-motor processing of tone changes in syllables was determined by language experience. This is the first study to provide causal evidence that the lateralization of motor contributions to auditory speech processing is determined by the functional role of the acoustic cues in the listener’s native language.

Our findings support the view that the left hemisphere selectively contributes to the processing of speech sounds when the auditory input is interpreted linguistically (14,23). We found that in tonal language speakers, disruption of the left motor cortex impaired auditory processing of both phoneme and tone changes. This is in line with early lesion studies with patients, reporting impaired lexical tone production and perception in tonal language speakers following damage to the left hemisphere (24-26) and dichotic listening studies, showing tonal language speakers could more accurately report lexical tones presented to their right ear (27,28). Later, functional neuroimaging data also revealed that tonal language speakers mainly recruit language regions in the left hemisphere for the processing of lexical tone (14-17, 29). For instance, compared with English speakers, Mandarin speakers exhibit greater tone-evoked activity in several regions in the left hemisphere, including the left posterior temporal and inferior parietal regions, known for their roles in auditory processing of language (15). Interestingly, the left ventral motor cortex has also been shown to be co-activated with auditory areas during discrimination of native tones (30). The current findings confirm that the left-hemisphere speech motor cortex affects auditory tone discrimination in tonal speakers.

In non-tonal language speakers, the disruption of the speech motor cortex in the right hemisphere impaired auditory tone discrimination. These results are consistent therefore with a known pattern of right hemisphere lateralization for tone and prosody perception based on findings from the lesion deficit (31), dichotic listening (32), and neuroimaging literature (10,11,33).

The differences in auditory-motor processing of tone changes in speech between the two language groups provide strong support for the influence of a listener’s language experience on hemispheric lateralization of speech. Lateralization of auditory speech processing is determined early during language acquisition: internationally adopted children who had been completely cut off from exposure to Mandarin since adoption (at 12.8 months of age, on average) exhibited left lateralization in the auditory system during lexical tone processing when studied over 10 years later as teenagers (17). Future studies are needed to investigate how lateralization of motor contributions to speech processing develops and is modulated by early and late language experience. In the current study, all native Mandarin speakers had a basic level of English and used both languages in their everyday life.

The motor contribution to auditory tone processing was strongly left-lateralized in tonal-language speakers, while it was weakly right-lateralized in non-tonal language speakers. Whereas the MMN responses elicited by tonal changes in tonal-language speakers were unchanged following right-hemisphere motor disruption, a small reduction in these responses in non-tonal language speakers was observed following left-hemisphere motor disruption. Such an effect was not observed in a previous study of non-tonal language speakers, in which behavioural discrimination of spoken questions and statements was impaired by disruption of right but not left premotor cortex (8). It is possible that the subtle effect of left-hemisphere disruption found in our study, which was considerably smaller than the effect of right-hemisphere disruption, was revealed by the use of the MMN as opposed to less sensitive behavioural measures of task performance. The MMN provides an automatic measure of tone discrimination removing any task demands including the need for attention. It is possible that the auditory-motor processing of tone is bilaterally organized with a weak right-hemisphere bias in non-tonal language speakers. However, when attention is focused on tones, auditory-motor processing in the right hemisphere becomes more prominent.

The current study confirms the contribution of speech motor cortex to auditory processing of phoneme and tone changes in both tonal and non-tonal language speakers. The findings therefore provide further evidence for the causal role of the speech motor cortex in speech processing (20,21,34). Contemporary neuroanatomical models of language processing propose that the motor and auditory systems are reciprocally connected (35). According to theory, during speech production, an efference copy is sent from the motor cortex to the auditory cortex to encode the expected auditory consequences of upcoming movements, while corrective commands are also sent from auditory cortex to motor cortex that drive compensatory speech movement (36,37). We suggest that this reciprocal connection between auditory and motor systems is also engaged during speech perception, enabling the motor system to contribute to auditory processing of speech sounds. Moreover, the complementary contribution of speech motor cortex to speech perception might be observed only when contextual information is not available, as in the current experimental setting (38); this could partially reconcile findings from studies in patients showing relatively intact speech comprehension after damage to brain areas of the frontal-motor network (39-41). In real life conditions, listeners, including patients with Broca’s aphasia can make use of contextual information to “comprehend” the words even when their speech perception is partially impaired.

The auditory-motor processing of speech sounds observed in the current study was not feature-specific: the disruption of speech motor cortex involved targeting the lip representation separately in each hemisphere. The disruption affected both phoneme and tone discrimination, even though tone production is controlled by the larynx. This is consistent with previous studies, which showed that disruption of the left lip representation suppressed MMN responses evoked by infrequent lip-related ‘ba’ and tongue-related ‘ga’ sounds that were presented among frequent tongue-related ‘da’ sounds (21,22). However, previous behavioural studies suggested that the contribution of the speech motor cortex in speech perception was somatotopically organized: TMS-induced disruption of the left lip representation only impaired discrimination of lip-related sounds (‘ba’ vs. ‘da’) leaving the ability to discriminate tongue-related sounds (‘ka’ vs. ‘ga’) unimpaired (19,42). The nonspecific effect observed in current study could be due to the proximity of the lip and larynx representations, which makes it impossible to disrupt the lip without affecting the adjacent larynx representation (43,44). An alternative explanation is that the feature-specificity of motor contributions to speech perception depends on attention. We showed previously that when speech sounds are ignored, the motor contributions to auditory speech processing are late and non-specific (>170 ms), whereas when speech sounds are attended, there is an additional early (<100 ms) motor effect on auditory speech processing, which is feature-specific (22).

Finally, we suggest that the asymmetry observed in the auditory system during speech perception might be modulated by the motor system. Recently, a modified version of the asymmetry-in-time sampling hypothesis has been proposed, in which the auditory processing is modulated by two different mechanisms (45,46): i) under non-speech listening conditions, auditory processing is supported by intrinsic auditory mechanisms, in line with acoustic hypotheses; ii) however, depending on individual brain differences or specific speech-listening conditions or both, the relative weight of top-down mechanisms can increase, which might affect the lateralization of auditory processing. During continuous speech perception, the speech-coupled oscillations in auditory cortex are significantly modulated by topdown signals from the frontal and motor cortex (47). In the current study, the modulatory effect of top-down signals from motor cortex on auditory processing was strongly left lateralized in tonal language speakers. This might partially account for left-lateralized auditory speech processing that is determined by linguistic experience and the functional role of acoustic cues.

To sum up, our study, for the first time, presents causal evidence that the contribution of the left and right speech motor cortex to auditory speech processing is determined by the functional role of acoustic cues in the listener’s native language. Such asymmetry observed in the motor system could modulate the asymmetry in the auditory system. Our study emphasizes the importance of understanding the interactions between bottom-up auditory mechanisms and top-down influences, when investigating hemispheric asymmetry of speech processing. We also call for more comparative studies between speakers of different languages, since ignoring the diversity of spoken languages can to lead to an incomplete understanding of the neurobiology of speech and language.

## Materials and Methods

### Participants

We recruited 32 adult human participants (16 male, 16 female) who were speakers of non-tonal languages and aged between 18 and 35 years and 32 tonal language (Mandarin) speakers (16 male, 16 female) aged between 18 and 34 years. None of the 64 participants had received any formal musical training that had lasted longer than 2 years. The native languages of the group of non-tonal language speakers were: English (21), French (2), Spanish (2), Czech (2), German (1), Italian (1), Romanian (1), Polish (1), and Dutch (1). None of the non-tonal language speakers had ever learned a tonal language but most had learned another language to varying levels of proficiency. All the tonal language speakers were native speakers of Mandarin and had at least a basic level of English; 29 were studying at an English-speaking university and all were living in the UK. The ages at which they first started to learn English ranged from 7 to 15 years old. Male and female participants in each language group were randomly assigned to receive either left or right hemisphere stimulation, so that groups were gender balanced. All participants reported no hearing problems and no personal or family history of seizures or other neurological disorders. The Central University (of Oxford) Research Ethics Committee approved the experimental protocol and participants gave informed consent. EEG Data from two non-tonal language speakers (both native English speakers; 1 male, 1 female) were excluded from the analyses because their mean mismatch negativity (MMN) amplitudes were extreme (greater than three standard deviations from the group mean MMN amplitude). The mean ages and gender balance for the four groups that contributed data to the analysis are as follows: non-tonal language, left hemisphere stimulation group: mean age 21.60 years, 8 males, 7 females; non-tonal language, right hemisphere stimulation group: mean age 22.60 years, 7 males, 8 females; tonal language, left hemisphere stimulation group: mean age 25.69 years, 8 males, 8 females ; tonal language, right hemisphere stimulation group: mean age 26.13 years, 8 males, 8 females . There was no significant difference between groups in terms of age.

### Procedure

For all participants, EEG was recorded during a 16.5-min oddball sequence were recorded during two sessions: a no-TMS baseline and a post-rTMS session. Session order was counterbalanced: half participants started with the no-TMS session followed by rTMS and then an immediate post-rTMS session, while the other half received rTMS first, the post-rTMS session immediately after, and then a no-TMS session at least 50 minutes later. TMS involved 15-min of low-frequency (0.6 Hz) sub-threshold repetitive stimulation over the lip representation in either the left or the right primary motor cortex. This protocol temporarily inhibits excitability in the target representation for a further 15 mins approximately (19).

The oddball sound sequence comprised infrequent tone and phoneme changes, which were expected to elicit mismatch negativity (MMN) responses (48-50). During no-TMS baseline sessions, event-related potentials (ERPs) were also recorded to two control sequences (see below). The order of presentation of oddball and control sequences was counter-balanced. Participants watched a silent movie and were instructed to ignore the sound sequences during all recordings.

### Stimuli

The oddball sequence contained a total of 1800 stimuli, including infrequent ‘ba1’ and ‘da4’ (probability = 0.1 for each) and frequent ‘da1’ syllables (probability = 0.8). Control sequences consisted of 400 repetitions each of the infrequent syllables in the oddball sequence: two control sequences, one for ‘ba1’ and one for ‘da4’. Syllables ‘da1’ (tone 1: high-level tone) and ‘da4’ (tone 4: high falling tone) shared the same phoneme but carried different tone contours, while ‘da1’ and ‘ba1’ had the same high-level tone (tone 1) but contained different phoneme information. Both tone and phoneme changes were expected to elicit MMNs. The original stimuli were produced by a female native Mandarin speaker. Using Praat software (Institute of Phonetic Sciences, University of Amsterdam, The Netherlands), we digitally normalized them to equal durations (150 ms) and intensity (70 db) and matched them for fundamental frequency (F0) for the first 35 ms of the syllable (see figure S1). The ‘da4’ syllable differed from the other two in the F0 trajectory after 35ms, which is clearly seen in Figure S1. The inter-stimulus interval was 400 ms in all sequences. Stimulus presentation was controlled via custom scripts in MATLAB (MathWorks).

### Transcranial Magnetic Stimulation

All TMS pulses were monophasic, generated by two Magstim 200s and delivered through a 70-mm figure-8 coil connected through a BiStim module (Magstim, Dyfed, UK).

Before rTMS, single TMS pulses were used to localize the lip representation, following the procedure previously described (51). Then, once the target area had been located, we established the active motor threshold, that is, the minimum intensity at which TMS elicited at least 5 out of 10 motor-evoked potentials (MEPs) with an amplitude of at least 200 μV when the target muscle was contracted at 20–30% of the maximum. The mean active motor threshold (percentage of maximum stimulator output, ±SD) for the left and right stimulation groups respectively was 63.1% (±7.8) and 62.1% (±7.1) in the tonal language speakers, and 58.8% (±6.1) and 57.47% (±7.4) in the non-tonal speakers. The intensity of each participant’s active motor threshold was used for 15-min rTMS while the muscles were relaxed (hence the stimulation was subthreshold). During rTMS, participants watched a silent nature documentary and were instructed to stay still. Electromyography (EMG) recordings were carefully monitored throughout stimulation to ensure that muscles were relaxed and no MEPs were evoked in the lip muscles. The coil was changed halfway through rTMS to prevent overheating.

### Electromyography (EMG) recordings

Disposable electrodes were attached on the right or left orbicularis oris muscle (the upper and lower lip contralateral to the stimulation site) and forehead (ground) to record electromyography (EMG) signals. The EMG signals were amplified, sampled at 5000 Hz, filtered (1 Hz–1 kHz bandpass) using a CED 1902 amplifier, a CED 1401 analog-to-digital converter, and a PC running Spike2 (Cambridge Electronic Design). A power bar displayed on a computer screen allowed participants to practise producing a constant level of contraction of the lips (20–30% of the maximum).

### Electroencephalographic (EEG) recording parameters

We acquired EEG using Synamps amplifiers and Curry 7 data acquisition software. We used a custom electrode cap with 11 electrodes (Fz, FCz, Cz, CPz, Pz, F1, F2, P1, P2, left, and right mastoids), which provided maximal coverage over sites of interest. The ground and reference electrodes were placed on the right upper arm and the tip of the nose, respectively. During acquisition, data were low-pass filtered (400 Hz cutoff), digitized at 1000 Hz, and stored for offline analysis. Electrode impedances were reduced below 10 kΩ before recording.

### EEG data Analyses

EEG signals were preprocessed using the EEGLab Toolbox (52). The raw data (down sampled to 500 Hz) were first re-referenced to the mean of the two mastoids to improve the signal-to-noise ratio of the MMN responses. Then the data were digitally filtered (low-pass filter of 30 Hz, high-pass filter of 1 Hz), baseline corrected and segmented into epochs of 400 ms comprising a 100-ms prestimulus baseline and a 300-ms interval after the onset of the stimuli. The epochs containing amplitude fluctuations exceeding ± 70 μV were considered as artifacts and removed before averaging. In addition, the epochs for the first 10 stimuli in each sequence and for the first standard stimulus after each deviant were removed.

MMN responses were calculated in two ways for the baseline (no TMS) sessions: the traditional method (subtracting the ERPs evoked by the frequent stimuli from those evoked by the infrequent stimuli in the oddball sequence), and the same-stimulus method (subtracting ERPs evoked by the stimuli presented frequently in the control sequence from those evoked by identical sounds presented infrequently in the oddball sequences). The advantage of the traditional method is that the responses are recorded for frequent and infrequent stimuli during the same sequence. The disadvantage is that we are comparing responses to sounds that differ acoustically. The same-stimulus method gets around this potential confound by comparing the response to acoustically identical stimuli presented either in the control sequence (i.e. with other identical stimuli) or in the oddball sequence as an infrequent sound (i.e. with other acoustically different stimuli). Identity MMN responses obtained using the same-stimulus method therefore reflect differences due to the automatic discrimination of speech sounds presented infrequently in the context of more frequent but acoustically different speech sounds (53-55). It should be noted that, due to the limited duration of the TMS-induced disruption (~15 min), control sequences were only recorded during the no-TMS session. Having confirmed that the MMN responses seen at baseline were due to discrimination responses (see Supplementary Tables S1 and S2), the traditional method was subsequently used to calculate the MMN responses in the post-rTMS sessions and compared with those obtained in the same way for the no-TMS sessions.

Statistical analyses followed the methods used in Möttönen, Dutton & Watkins (21). The FCz electrode was chosen as MMN responses are maximal over the fronto-central area (48,49). First, no-TMS and post-rTMS MMN responses were compared at each time point from 0 to 300 ms after the beginning of the syllable, to determine the latency at which there was a significant difference between no-TMS and post-rTMS MMN responses. The MMN differences were considered as significant when p-values were lower than 0.05 for 10 (= 20 ms) or more consecutive time points (21,56). The MMN amplitudes were calculated by averaging the responses across a 40-ms window centred at the peak of MMN. We then compared mean amplitudes of MMN responses elicited in the oddball sequence during no-TMS and post-rTMS recordings, and identity MMN calculated by the same-stimulus method (see above), with zero respectively with one-sample tests (two-tailed) to test whether significant MMN responses were evoked during different time points. To further test the influence of language experience on the lateralization of auditory-motor processing of speech sound, repeated measures analyses of variance (ANOVAs) were conducted separately for MMN responses elicited by phoneme and tone contrasts, with TMS (no-TMS vs. post-rTMS) as a within-subjects factor and language group (tonal language speakers v.s. non-tonal language speakers) as a between-subjects factor. Paired t-tests were then used (no-TMS vs. post-rTMS) in postdoc comparisons.

## Data availability

The raw EEG data generated in this study have been deposited in https://osf.io/en7uq/ The data information is summarised in a word document available on OSF.

## Acknowledgments

This research was supported by a studentship to D-L.T. from the China Scholarship Council. The Wellcome Centre for Integrative Neuroimaging is supported by core funding from the Wellcome Trust (203139/Z/16/Z). We thank Professor Kia Nobre for access to EEG facilities, Mark Roberts for technical support, and Wen Wen for EEG data collection.

## Author Contributions

D-L. T., K.E.W., and R.M. designed the experiments. D-L. T and S.S. A. recruited and tested participants. D-L. T analysed the data, created the figures and drafted the manuscript. All authors edited the manuscript for publication.

## Supplemental Information

Supplemental Information includes one figure and two tables

**Fig. S1.**
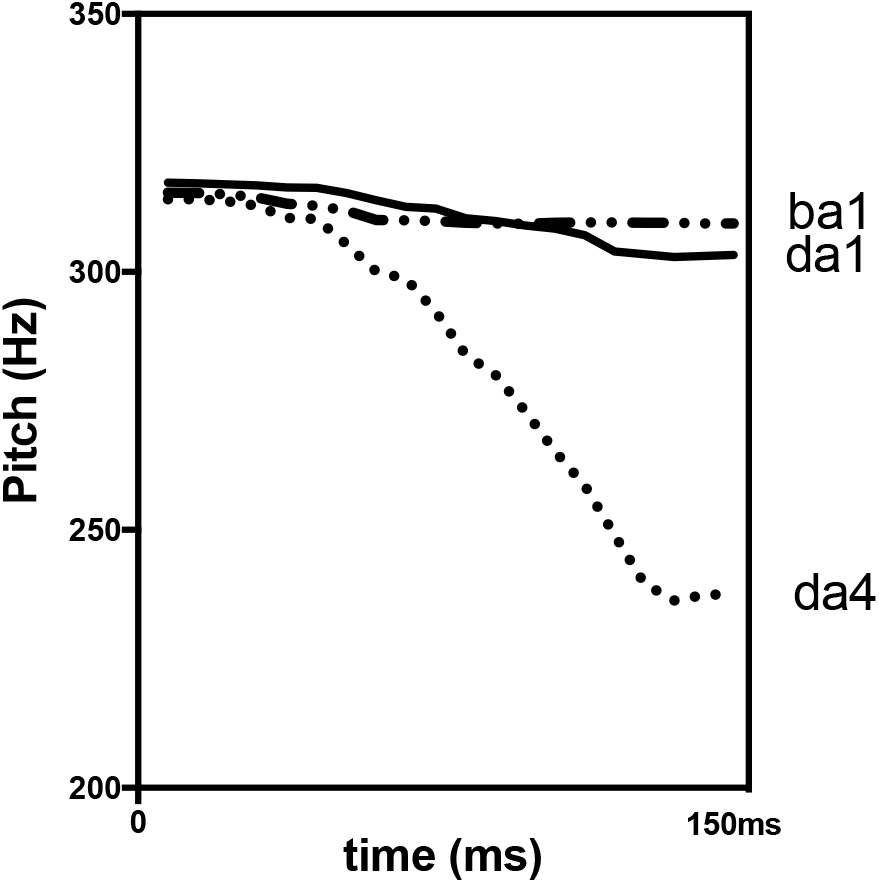
The trajectories of fundamental frequency (F0) of the three syllables used in current study. Significant identity MMN responses, calculated using the same-stimulus method, were observed during the no-TMS recording in all the four groups, confirming that MMN responses were elicited by automatic discrimination of speech sounds, rather than the acoustic differences between them (Table S1, S2).

**Table S1.** Left M1 stimulation: *Identity* MMN peak latencies (ms) and mean amplitudes (μV, ±SEM) at FCz.

|  | Peak | Amplitude | t |
| --- | --- | --- | --- |
| **Non-tonal language speakers, n=15** |  |  |  |
| 'ba1' | 210.4 | -2.20(0.30) | 7.41*** |
| 'da4' | 188.8 | -7.24(0.71) | 10.21*** |
| **Tonal language speakers, n=16** |  |  |  |
| 'ba1' | 204.6 | -2.64(0.41) | 6.34*** |
| 'da4' | 193.3 | -7.62(0.68) | 11.28*** |

Note: The identity MMN responses were calculated by subtracting event-related potentials evoked by the stimuli presented frequently (probability=1.0) in the control sequence from those evoked by identical sounds presented infrequently (probability= 0.1) in the oddball sequences (with no TMS). Mean amplitudes were calculated by averaging the responses across a 40-ms window centred at this peak latency and were compared to zero with 2-tailed t-tests.
\*p < .05
\*\*\*p < .001

**Table S2.**
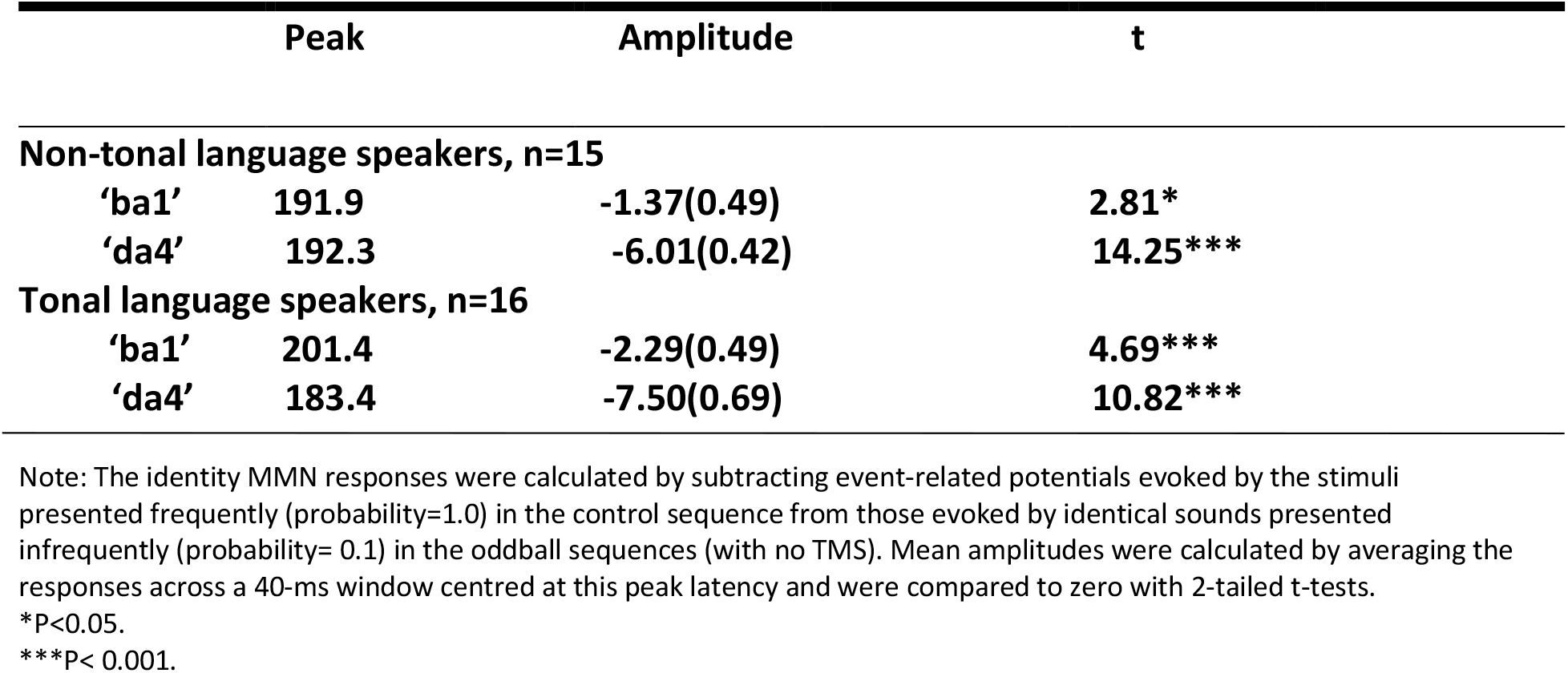
Right M1 stimulation: *Identity* MMN peak latencies (ms) and mean amplitudes (μV, ±SEM) at FCz.

## References

1. Skipper, J.I., Devlin, J.T., and Lametti, D.R. (2017). The hearing ear is always found close to the speaking tongue : Review of the role of the motor system in speech perception. Brain Lang. 164, 77–105.

2. Liebenthal, E., and Möttönen, R. (2018). An interactive model of auditory-motor speech perception. Brain Lang. 187, 33–40.

3. Fadiga L, Craighero L, Buccino G, Rizzolatti G. (2002). Speech listening specifically modulates the excitability of tongue muscles, a TMS study. Eur J Neurosci. 15:399–402.

4. Meister IG, Wilson SM, Deblieck C, Wu AD, Iacoboni M. (2007). The essential role of premotor cortex in speech perception. Curr Biol. 17:1692–1696.

5. Hickok, G., and Poeppel, D. (2007). The cortical organization of speech processing. Nat. Rev. Neurosci. 8, 393–402.

6. Wildgruber, D., Ackermann, H., Kreifelts, B., and Ethofer, T. (2006). Cerebral processing of linguistic and emotional prosody: fMRI studies. Prog. Brain Res. 156, 249–268.

7. Ethofer, T., Anders, S., Erb, M., Herbert, C., Wiethoff, S., Kissler, J., Grodd, W., and Wildgruber, D. (2006). Cerebral pathways in processing of affective prosody: a dynamic causal modeling study. Neuroimage 30, 580–587.

8. Sammler, D., Grosbras, M.-H., Anwander, A., Bestelmeyer, P.E.G., and Belin, P. (2015). Dorsal and Ventral Pathways for Prosody. Curr. Biol. 25, 3079–3085.

9. Zatorre, R.J., and Belin, P. (2001). Spectral and Temporal Processing in Human Auditory Cortex. Cereb. Cortex 11, 946–953.

10. Boemio, A., Fromm, S., Braun, A., and Poeppel, D. (2005). Hierarchical and asymmetric temporal sensitivity in human auditory cortices. Nat. Neurosci. 8, 389–395.

11. Albouy, P., Benjamin, L., Morillon, B., and Zatorre, R.J. (2020). Distinct sensitivity to spectrotemporal modulation supports brain asymmetry for speech and melody. Science (80-.). 367, 1043–1047.

12. Jamison, H. L., Watkins, K. E., Bishop, D. V., & Matthews, P. M. (2006). Hemispheric specialization for processing auditory nonspeech stimuli. Cereb Cortex, 16(9), 1266–1275.

13. Yip, M., 2002. Introduction. Tone. Cambridge University Press, Cambridge, England 1–16. Yost, W.A., 2009. Pitch perception. Atten. Percept. Psychophys. 71, 1701–1715.

14. Hsieh, L., Gandour, J., Wong, D., and Hutchins, G.D. (2001). Functional Heterogeneity of Inferior Frontal Gyrus Is Shaped by Linguistic Experience. Brain Lang. 76, 227–252.

15. Klein, D., Zatorre, R.J., Milner, B., and Zhao, V. (2001). A Cross-Linguistic PET Study of Tone Perception in Mandarin Chinese and English Speakers. Neuroimage 13, 646–653.

16. Gandour, J., Wong, D., Hsieh, L., Weinzapfel, B., Lancker, D. Van, and Hutchins, G.D. (2000). A Crosslinguistic PET Study of Tone Perception. J. Cogn. Neurosci. 12, 207–222.

17. Pierce, L.J., Klein, D., Chen, J.-K., Delcenserie, A., and Genesee, F. (2014). Mapping the unconscious maintenance of a lost first language. Proc. Natl. Acad. Sci. 111, 17314–17319.

18. Zatorre, R. J. & Gandour, J. T. (2008). Neural specializations for speech and pitch: moving beyond the dichotomies. Philos. Trans. R. Soc. Lond., B, Biol. Sci. 363, 1087–1104.

19. Mottonen, R., and Watkins, K.E. (2009). Motor Representations of Articulators Contribute to Categorical Perception of Speech Sounds. J. Neurosci. 29, 9819–9825.

20. Smalle, E.H.M., Rogers, J., and Möttönen, R. (2015). Dissociating Contributions of the Motor Cortex to Speech Perception and Response Bias by Using Transcranial Magnetic Stimulation. Cereb. Cortex 25, 3690–3698.

21. Mottonen, R., Dutton, R., and Watkins, K.E. (2013). Auditory-Motor Processing of Speech Sounds. Cereb. Cortex 23, 1190–1197.

22. Mottonen, R., van de Ven, G.M., and Watkins, K.E. (2014). Attention Fine-Tunes Auditory-Motor Processing of Speech Sounds. J. Neurosci. 34, 4064–4069.

23. Liberman, A.M., and Whalen, D.H. (2000). On the relation of speech to language. Trends Cogn. Sci. 4, 187–196.

24. Gandour, J., Ponglorpisit, S., and Dardarananda, R. (1992). Tonal disturbances in Thai after brain damage. J. Neurolinguistics 7, 133–145.

25. Yiu, E.M.-L., and Fok, A.Y.-Y. (1995). Lexical tone disruption in Cantonese aphasic speakers. Clin. Linguist. Phon. 9, 79–92.

26. Eng, N., Obler, L.K., Harris, K.S., and Abramson, A.S. (1996). Tone perception deficits in Chinese-speaking Broca’s aphasics. Aphasiology 10, 649–656.

27. Van Lancker, D., and Fromkin, V.A. (1973). Hemispheric specialization for pitch and “tone”: Evidence from Thai. J. Phon. 1, 101–109.

28. Van Lancker, D. (1980). Cerebral lateralization of pitch cues in the linguistic signal. Pap. Linguist. 13, 201–277.

29. Gandour, J., Dzemidzic, M., Wong, D., Lowe, M., Tong, Y., Hsieh, L., Satthamnuwong, N., and Lurito, J. (2003). Temporal integration of speech prosody is shaped by language experience: An fMRI study. Brain Lang. 84, 318–336.

30. Xu, Y., Gandour, J., Talavage, T., Wong, D., Dzemidzic, M., Tong, Y., Li, X., and Lowe, M. (2006). Activation of the left planum temporale in pitch processing is shaped by language experience. Hum. Brain Mapp. 27, 173–183.

31. Zatorre, R.J., and Samson, S. (1991). Role of the right temporal neocortex in retention of pitch in auditory short-term memory. Brain. 114, 2403–2417.

32. Sidtis, J.J. (1980). On the nature of the cortical function underlying right hemisphere auditory perception. Neuropsychologia 18, 321–330.

33. Schönwiesner, M., Rübsamen, R., and Von Cramon, D.Y. (2005). Hemispheric asymmetry for spectral and temporal processing in the human antero-lateral auditory belt cortex. Eur. J. Neurosci. 22, 1521–1528.

34. Davis, M.H., and Johnsrude, I.S. (2007). Hearing speech sounds: Top-down influences on the interface between audition and speech perception. Hear. Res. 229, 132–147.

35. Pulvermüller, F., and Fadiga, L. (2010). Active perception: sensorimotor circuits as a cortical basis for language. Nat. Rev. Neurosci. 11, 351–360.

36. Tourville, J.A., and Guenther, F.H. (2011). The DIVA model: A neural theory of speech acquisition and production. Lang. Cogn. Process. 26, 952–981.

37. Houde, J.F., and Nagarajan, S.S. (2011). Speech Production as State Feedback Control. Front. Hum. Neurosci. 5.

38. Wilson, S.M. (2009). Speech perception when the motor system is compromised. Trends Cogn. Sci. 13, 329–330.

39. Craighero, L., Metta, G., Sandini, G., and Fadiga, L. (2007). The mirror-neurons system: data and models. In Progress in Brain Research, pp. 39–59.

40. Hickok, G., Okada, K., Barr, W., PA, J., Rogalsky, C., Donnelly, K., Barde, L., and Grant, A. (2008). Bilateral capacity for speech sound processing in auditory comprehension: Evidence from Wada procedures. Brain Lang. 107, 179–184.

41. Hickok, G. (2010). The role of mirror neurons in speech perception and action word semantics. Lang. Cogn. Process. 25, 749–776.

42. D’Ausilio, A., Pulvermüller, F., Salmas, P., Bufalari, I., Begliomini, C., and Fadiga, L. (2009). The Motor Somatotopy of Speech Perception. Curr. Biol. 19, 381–385.

43. Grabski, K., Lamalle, L., Vilain, C., Schwartz, J.-L., Vallée, N., Tropres, I., Baciu, M., Le Bas, J.-F., and Sato, M. (2012). Functional MRI assessment of orofacial articulators: Neural correlates of lip, jaw, larynx, and tongue movements. Hum. Brain Mapp. 33, 2306–2321.

44. Eichert. N., Papp. D., Mars. R.B., Watkins, K.E. (2020). Mapping human laryngeal motor cortex during vocalization. BioRxiv.

45. Assaneo, M. F., Rimmele, J. M., Orpella, J., Ripolles, P., de Diego-Balaguer, R., & Poeppel, D. (2019). The Lateralization of Speech-Brain Coupling Is Differentially Modulated by Intrinsic Auditory and Top-Down Mechanisms. Front Integr Neurosci, 13, 28.

46. Rimmele, J. M., Morillon, B., Poeppel, D., and Arnal, L. H. (2018). Proactive sensing of periodic and aperiodic auditory patterns. Trends Cog. Sci. 22, 870–882.

47. Park, H., Ince, R. A., Schyns, P. G., Thut, G., & Gross, J. (2015). Frontal top-down signals increase coupling of auditory low-frequency oscillations to continuous speech in human listeners. Curr Biol, 25(12), 1649–1653. doi:10.1016/j.cub.2015.04.049.

48. Näätänen, R., Lehtokoski, A., Lennes, M., Cheour, M., Huotilainen, M., Iivonen, A., Vainio, M., Alku, P., Ilmoniemi, R.J., Luuk, A., et al. (1997). Language-specific phoneme representations revealed by electric and magnetic brain responses. Nature 385, 432–434.

49. Näätänen, R., Tervaniemi, M., Sussman, E., Paavilainen, P., and Winkler, I. (2001). ‘Primitive intelligence’ in the auditory cortex. Trends Neurosci. 24, 283–288.

50. Winkler, I., Kujala, T., Tiitinen, H., Sivonen, P., Alku, P., Lehtokoski, A., Czigler, I., Csépe, V., Ilmoniemi, R.J., and Näätänen, R. (1999). Brain responses reveal the learning of foreign language phonemes. Psychophysiology 36, S0048577299981908.

51. Möttönen, R., Rogers, J., and Watkins, K.E. (2014). Stimulating the Lip Motor Cortex with Transcranial Magnetic Stimulation. J. Vis. Exp.

52. Delorme, A., and Makeig, S. (2004). EEGLAB: an open source toolbox for analysis of single-trial EEG dynamics including independent component analysis. J. Neurosci. Methods 134, 9–21.

53. Jacobsen, T., and Schröger, E. (2001). Is there pre-attentive memory-based comparison of pitch? Psychophysiology 38, S0048577201000993.

54. Jacobsen, T., and Schröger, E. (2003). Measuring duration mismatch negativity. Clin. Neurophysiol. 114, 1133–1143.

55. Kujala, T., Tervaniemi, M., and Schröger, E. (2007). The mismatch negativity in cognitive and clinical neuroscience: Theoretical and methodological considerations. Biol. Psychol. 74, 1–19.

56. Guthrie, D., and Buchwald, J.S. (1991). Significance Testing of Difference Potentials. Psychophysiology 28, 240–244.

